# High pressure inhibits signaling protein binding to the flagellar motor and bacterial chemotaxis through enhanced hydration

**DOI:** 10.1101/762922

**Authors:** Hiroaki Hata, Yasutaka Nishihara, Masayoshi Nishiyama, Yoshiyuki Sowa, Ikuro Kawagishi, Akio Kitao

## Abstract

In the chemotaxis of *Escherichia coli*, the cell’s behavioral switch involves binding of the phosphorylated form of the chemotaxis signaling protein CheY (CheYp) to the flagellar motor protein FliM, which induces the motor to rotate clockwise; otherwise, the motor rotates counterclockwise. To investigate high-pressure effects on CheYp–FliM binding at atomic resolution, we conduct molecular dynamics simulations of monomeric CheYp, the N-terminal fragment of the FliM (FliM_N_) that binds to CheYp, and the complex that forms between those proteins at pressures ranging from 0.1 to 100 MPa. The results show that the active form of monomeric CheYp is maintained even at 100 MPa but high pressure increases the water density in the first hydration shell and can cause conformational change of the C-terminal helix. The dissociation process of the complex is investigated by parallel cascade selection molecular dynamics (PaCS-MD), revealing that high pressure considerably induces water penetration into the complex interface. Pressure dependence of standard binding free energy calculated by the Markov state model indicates that the increase of pressure from 0.1 to 100 MPa weakens the binding by ∼ 10 kcal/mol. Using high-pressure microscopy, we observed that high hydrostatic pressure reversibly fixes the motor rotation in the counter-clockwise orientation, which supports the notion that high pressure inhibits the binding of CheYp to FliM. We conclude that high pressure induces water penetration into the complex interface, which interferes with CheYp–FliM binding and prevents motor reversal.

## Introduction

Pressure significantly affects protein structure, dynamics, and functions^1–4^. Very high pressure (above 500 MPa) induces denaturation of many soluble proteins^5–8^, and therefore the mechanism of pressure denaturation has been relatively well investigated, both theoretically and experimentally^4,9,10^. On the other hand, pressures below 100 MPa, a range that covers the typical biosphere on earth, including the deep sea, does not induce large changes in overall secondary and tertiary structure of most protein molecules^1,3,11,12^, but was found to affect behaviors of biological systems at macroscopic levels^13–15^. Pressures of ∼ 100 MPa are known to interfere with interactions between biomolecules such as proteins and ligands^1,4,14,16,17^, indicating that the mechanism whereby high pressure perturbs such interactions should be elucidated at the molecular level.

Molecular dynamics (MD) simulation has been applied to obtain atomic details of structures and functions of biomolecules, including surrounding water and other molecules^18–20^. MD simulations under high pressures have observed enhanced hydration of proteins^21–23^ and penetration of water molecules inside proteins with accompanying conformational changes^24,25^. However, it remains difficult to simulate pressure effects on protein-protein interactions, given that the timescale for observing the pressure effects is often longer than the length of typical MD simulations^4,26^. Recently, a new distributed computing method, parallel cascade selection molecular dynamics (PaCS-MD), has been developed to observe events whose timescales are longer than the standard MD length^27,28^. PaCS-MD can simulate dissociation process of protein complexes, whose time scales range from μs to s, without using artificial forces within a short MD simulation time^29^. Additionally, by integrating the distributed MD trajectories using a Markov state model (MSM)^30^, various quantities characterizing molecular interactions can be calculated, such as binding free energy, association/dissociation rate constants, and residence times^29,31^.

In the present study, we investigate the effect of high pressure on the interactions between the chemotaxis signaling protein CheY and the flagellar rotor protein FliM by a combination of MD, PaCS-MD/MSM, and high-pressure microscopy. Switching of the bacterial flagellar motor from counter-clockwise (CCW) to clockwise (CW) is triggered by the binding of the phosphorylated CheY (CheYp) to the N-terminal segment of FliM (FliM_N_)^32,33^, which is essential for bacterial chemotaxis^34^. During CCW rotation, multiple flagellar filaments form a bundle, which smoothly propels the bacteria; in contrast, CW rotation untangles the bundle and consequently leads to changes in the swimming direction. High-pressure microscopy has revealed various phenomena induced in the flagellar motor by pressure^15,35^. *Escherichia coli* cells stop swimming at 80 MPa even if the flagellar motors generate sufficient torque, which implies the inhibition of flagellar bundle formation^35^. Application of pressure >120 MPa induces a reversal from CCW to CW in the absence of CheYp, suggesting pressure-induced structural changes similar to those caused by the binding of CheY^15^. At the molecular level, however, high-pressure effects on the CheYp–FliM binding have not been fully explored.

To investigate the CheYp–FliM interaction, we first examine the protein structure of CheY and FliM_N_ at pressures up to 100 MPa by standard MD simulations, which identify changes in the protein structures and the hydration state induced by high pressure. We then use PaCS-MD to simulate the high-pressure dissociation process of the CheYp-FliM_N_ complex, demonstrating that pressure increased hydrated waters and enhanced penetration of water molecules into the complex interface. MSM analysis quantitatively shows that high pressure decreases the binding free energy between CheYp and FliM_N_. This tendency is consistent with the microscopy microscopic observations, which showed that high hydrostatic pressure exclusively fixes the motor rotation in the CCW orientation at 40 MPa. We conclude that the application of pressure enhances hydration of the proteins and weakens the binding of CheYp to FliM_N_, resulting in CCW rotation of the flagellar motor.

## Methods

### MD simulation of monomeric CheY, FliM_N_, and the complex

First, pressure effects on monomeric CheY, monomeric FilM_N_, and their complex were investigated by standard MD simulation. We prepared four different models of CheY (aCheYp, iCheYp, aCheY, and iCheY). The structure of aCheY was generated by removing FliM_N_ from the aCheY-FliM_N_ complex structure (PDB ID: 1F4V^36^). The monomeric CheY crystal structure (PDB ID: 3CHY^37^) was used for iCheY. The aCheYp and iCheYp structures were generated by modeling the phosphorylated Asp57 residue based on the crystal structure of an *α*-thiophosphonate derivative of the CheY D57C point mutant (PDB ID: 1C4W^38^). The N-terminal fragment (residues 1–16) of the monomeric FliM_N_ was obtained by removing aCheY from the crystal structure of the complex. The crystal structure of BeF_3_^−^-activated CheY in complex with FliM_N_ comprising residues 1–16 shows that the first 7 residues of FliM_N_ takes an extended conformation and residues 8–15 forms an α-helix^36^. The CheYp–FliM_N_ complex structure was modeled based on the crystal structure (PDB ID: 1F4V^36^) by replacing BeF_3_^−^ with a phosphate group.

After energy minimization, each of the above protein models was solvated into a cubic periodic box and gaps were filled with water molecules and 0.15 M KCl. The prepared systems contain a total of ∼ 34,000, 23,000 and 38,000 atoms for monomeric CheY, monomeric FliM_N_, and the CheYp–FliM_N_ complex, respectively. All of the MD simulations were performed using the GPU implementation^39^ of the PMEMD module of the Amber14 package. The AMBER ff14SB force field^40^ was used for the proteins and ions. The TIP3P model^41^ was used for water molecules. The atomic partial charges of the phosphorylated aspartate were determined by quantum calculation using GAMESS^42^ and the RESP^43^ charge calculation. The other parameters (bond, angle, and dihedral) of the atoms were obtained using the general Amber force field (GAFF)^44^. These parameter files were generated by the module Antechamber of the AMBER package. The covalent bonds involving hydrogens were constrained using the SHAKE algorithm^45^, and the water molecules were kept rigid using the SETTLE algorithm^46^. After a 4000-step energy minimization, the system was brought to thermodynamic equilibrium at 300 K and 0.1, 50, or 100 MPa, using a Langevin thermostat^47^ and a Monte Carlo barostat. Equations of motion were integrated with a time step of 2 fs. The long-range Coulomb energy was evaluated using the particle mesh Ewald method^48^. The MD simulations were conducted for 0.2, 0.5, and 1.0 *μ*s for monomeric CheY, monomeric FliM_N_, and the complex of CheYp and FliM_N_, respectively. Two pressure conditions of 0.1 and 100 MPa were used for the simulations of monomeric CheY, and three conditions (0.1, 50, and 100 MPa) were employed for monomeric FliM_N_ and the CheYp–FliM_N_ complex.

The solvent accessible surface area (SASA) and the excluded volume (*V*_*ex*_; defined as the inside space of SASA) for CheY, FliM_N_, and the CheYp–FliM_N_ complex were calculated using the CAVE software package^49^. The calculation of SASA for the interface residues of the CheYp–FliM_N_ complex and the other analyses were performed using the cpptraj module of the AmberTools 14 Package. The representative MD structures were determined by k-means clustering into five clusters from the last half of each MD trajectory. Secondary structure was assigned by the DSSP program^50^. The isothermal compressibility, *κ*_*T*_, was obtained as follows^11^.

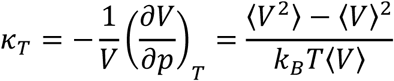

where *V, p, k*_*B*_, and *T* represent the system volume, pressure, the Boltzmann constant, and the absolute temperature, respectively. The angle bracket here denotes the average over the last half of the simulation. To calculate the density of bulk water, MD simulations with 10,056 water molecules were performed at 300 K and 0.1, 50, and 100 MPa for 100 ns. The average density during the last 50 ns of the MD trajectories were described in the text as the bulk water density.

### Dissociation simulation of the CheYp–FliM_N_ complex by PaCS-MD

To observe the dissociation process of the aCheYp and FliM_N_ complex, we used a PaCS-MD simulation^27,28^ in which mutually-overlapping conformational trajectories are generated by cycles of parallel short MD simulations and selection of MD snapshots closer to dissociation without applying additional bias to the system. In earlier works, PaCS-MD was successful in generating natural dissociation pathways of tri-N-acetyl-d-glucosamine from hen egg white lysozyme^31^ and those of the transactivation domain of the p53 protein from the MDM2 protein^29^. In the present work, the dissociation process of the complex was investigated by a procedure similar to that used in the earlier reports.

As the initial structures for PaCS-MD, the structures of the CheYp–FliM_N_ complex after 200 ns MD at 0.1, 50, and 100 MPa were adopted. To observe the dissociation up to *d* = 50 *Å*, the simulation box was expanded to a cubic box of 83 × 83 × 83 *Å*^*3*^ and the gaps were filled with water and 0.15 M KCl. The total numbers of atoms in the systems were ∼ 56,000 atoms including water and ions. After a short energy minimization, the simulation system was equilibrated at 300 K and 0.1 (or 50 or 100) MPa for 10 ns. The dissociation simulation was conducted five times with different initial conformations selected every 1 ns from the last snapshot of the 10-ns MD trajectory. For each PaCS-MD cycle, ten 100-ps MD simulations were performed in parallel and the trajectories were recorded every 10 fs. The snapshots sampled every 1 ps were rank-ordered according to the inter-COM distance, *d*, between CheYp and the helical segment of FliM_N_ (residues 8–16), and the top ten snapshots were selected for the next cycle. The first 7 residues of FliM_N_ were excluded from the calculation of COM because we noticed in preliminary trials that this region is very flexible and its fluctuation creates ‘noise’ in detecting the dissociation; the inclusion of the flexible region made detection of the dissociative movements more difficult. This restriction was applied only in the selection in PaCS-MD; all residues of FliM_N_ were considered for the calculation of the inter-COM distances in other determinations. No positional restraints were applied, thereby allowing reorientation of CheYp. In the present work, we used a larger cubic box so that FliM_N_ was permitted to dissociate in any direction. This approach differed from that used in the earlier work, in which reorientation of a protein was restrained within a rectangular box [12].

### Calculation of binding free energy by MSM

Standard binding free energies of CheYp and FliM_N_ at pressures of 0.1, 50, and 100 MPa were calculated using the MSM analysis of the MD trajectories^30^ generated by each trial. For a reasonable estimation of the transition probabilities among microstates in MSM, the trajectories from 10 independent MD simulations typically were not sufficient to achieve meaningful statistics. Instead, 10 additional MD simulations were performed, starting from the initial structures of each cycle of the PaCS-MD but with different initial velocities. This addition was repeated until sufficient statistics was achieved according to the implied time scale test^30,51^. For the 0.1 MPa condition, one set of the additional MD simulations (10 MDs) was performed for two of the five PaCS-MD trials, and two sets (20 MDs) were performed for the other three trials. At 50 MPa, one additional set (10 MDs) was performed for two trials, three sets (30 MDs) were performed for one trial, and the other two trials were analyzed by MSM without additional MD simulations. At 100 MPa, one additional MD set was performed for four trials, and two sets were performed for the rest.

Clustering and MSM construction were performed for each dissociation simulation using MSMBuilder 3.5.0^52^. The inter-COM distance between CheYp and FliM_N_ were calculated for all of the snapshots and were clustered into 30 clusters based on the k-means. The same procedure previously was used successfully in reproducing experimentally measured standard binding free energy of a protein-peptide complex^29^. A lag time of 30 ps was selected based on the implied time scales plot as a function of lag time. Free energy profiles (potential of mean force) were first calculated from the stationary probabilities of the microstates of each trial and then averaged over the multiple trials. The bound state was defined as a region before the free energy curve became flat, i.e., 40, 40, and 25 Å of the inter-COM distance for 0.1, 50 and 100 MPa, respectively. The other region (i.e., up to 50 Å after the energy curve became flat) was defined as the unbound state. The free energy difference (*ΔG*bind) was calculated from the probabilities of the bound (*P*_b_) and unbound (*P*_u_) states:

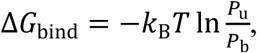

where *k*_*B*_ is the Boltzmann constant and *T* is the absolute temperature. The values of *P*_b_ and *P*_u_ were obtained by summing the probabilities of the microstates included in the respective state. The standard free energy of binding 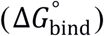 was calculated using the following expression with a volume correction term^53^:

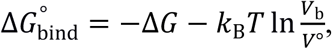

where *V*_b_ is the bound volume and *V*° is the standard-state volume (1661 *Å*^*3*^). The value of *V*_b_ was calculated as the volume of the convex hull defined by the COM coordinates of FliM_N_ in the bound state relative to the COM of CheYp at the origin. The convex hull calculation was performed for each dissociation simulation using Qhull^54^. Snapshots sampled every 2 ps were used for the *V*_b_ calculation. The averaged value of *V*_b_ for each pressure condition was used for the *ΔG*° calculation.

### Motility assay

Bacterial assays were performed using *E. coli* strain SYC12 (*fliC-sticky*, wild type for chemotaxis), a derivative of RP437^55^. SYC12 carries the *fliC*^*st*^ allele, encoding a ‘sticky’ mutant FliC filament protein that lacks residues 245-301 of what would otherwise be a 497-residue protein^56^; the mutant protein polymerizes into a filament that readily adheres to hydrophobic surfaces^57^. Following growth, the cells were resuspended in MLM medium (10 mM potassium phosphate (pH 7.0), 0.1 mM EDTA, 10 mM DL-lactate, 0.01 mM methionine) as described previously^58^. The motor rotation at ambient pressure and 40 MPa was recorded at 30 frames per second by high-pressure microscopy^15,35,59^. The pressure was controlled with an accuracy of ±1 MPa. The experimental temperature was maintained at 23 ± 1°C. After the release of pressure, all cells were removed from the chamber, and the next assay was performed using naïve cells (i.e., cells that had not been exposed previously to high pressure). All assays were repeated with at least three different cultures.

## Results and Discussion

### CheYp active form is stable even at high pressure

First, we examined the pressure effects on phosphorylation-dependent stability of monomeric CheY active form. Phosphorylation of CheY at the Asp57 sidechain induces a conformational change of CheY from the inactive (Fig. 1, orange) to the active (Fig. 1, green) form that is mainly characterized by a transition of the sidechain *χ*1 angle of Tyr106 from ∼ 60° to ∼ −150°^36,37^, resulting in an increase in the binding affinity of CheY for the N-terminal segment of FliM (FliM_N_)^36,60^. To model the combination of the active (Tyr106 *χ*1 ∼ 60°) or inactive (∼ −150°) form, phosphorylated (CheY) or non-phosphorylated at Asp57 (CheYp), four different models of CheY were constructed and simulated for 1 μs by MD at 0.1 and 100 MPa.

**Figure 1.**
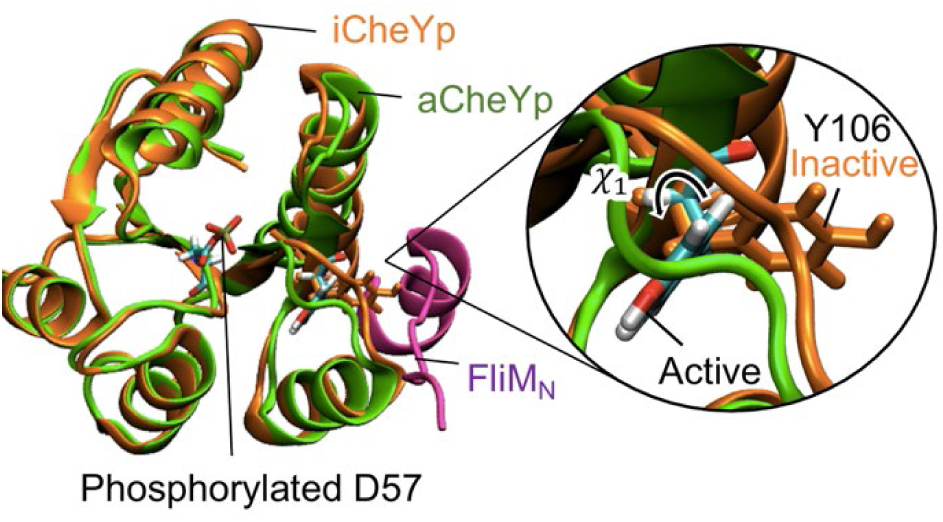
Molecular structures of CheYp and FliM_N_. Initial MD structures of the phosphorylated CheY in the active (aCheYp: green) and inactive (iCheYp: orange) forms. Y106 and the phosphorylated D57 residues are shown in stick models and those of aCheYp are colored on an atomic-color basis. A close-up view of Y106 is shown on the right. The FliM_N_ structure complexed with aCheYp is shown in magenta. In this paper, the molecular structure was visualized using VMD^75^.

In the MD simulations of CheYp started from the active form (aCheYp: *χ*_1_ = −150°), *χ*_1_ remained at approximately −150° at both 0.1 and 100 MPa (upper panel in Fig. 2A). Even when the MD of CheYp was started from the inactive form (iCheYp: *χ*_1_ = 60°), *χ*_1_ angle made a transition from 60° to −150° and remained at approximately −150° at both pressure conditions (second upper panel in Fig. 2A). These results clearly indicated that, for CheYp, the active form is more favorable than the inactive form in the pressure range ≤ 100 MPa. Also, the observed stabilization of phospho-CheY in the active form at ambient pressure is consistent with previous experimental observations^38,61,62^. It is worth mentioning that the time scale of the active-inactive transition is slowed at 100 MPa. In the case of iCheYp, the transition at 100 MPa occurred after 120 ns, while this transition was observed within 20 ns at 0.1 MPa. The same trend was also seen in the other cases. This slowing presumably reflects the pressure-induced slowdown of molecular motion and a consequent stiffening^63–65^. In the case of non-phosphorylated CheY, the active form is more destabilized. Non-phosphorylated CheY (lower two panels of Fig. 2A) tended to make conformational transitions between the active and inactive forms, with the exception of iCheY at 0.1 MPa. Starting from the active form without phosphorylation (aCheY) at 0.1 MPa, CheY exhibited conformational transitions between the two forms and eventually shifted predominantly to the inactive form, while at 100 MPa, CheY mostly remained in the active form; this pattern presumably resulted the slowed conformational change at high pressure. MD of non-phosphorylated CheY started from the inactive (iCheY) underwent no such transition and retained the inactive form at 0.1 MPa; few transitions were observed at 100 MPa. Overall, the MD simulation of monomeric CheY clearly indicated that activated phosho-CheY is not affected by high pressure below 100 MPa.

**Figure 2.**
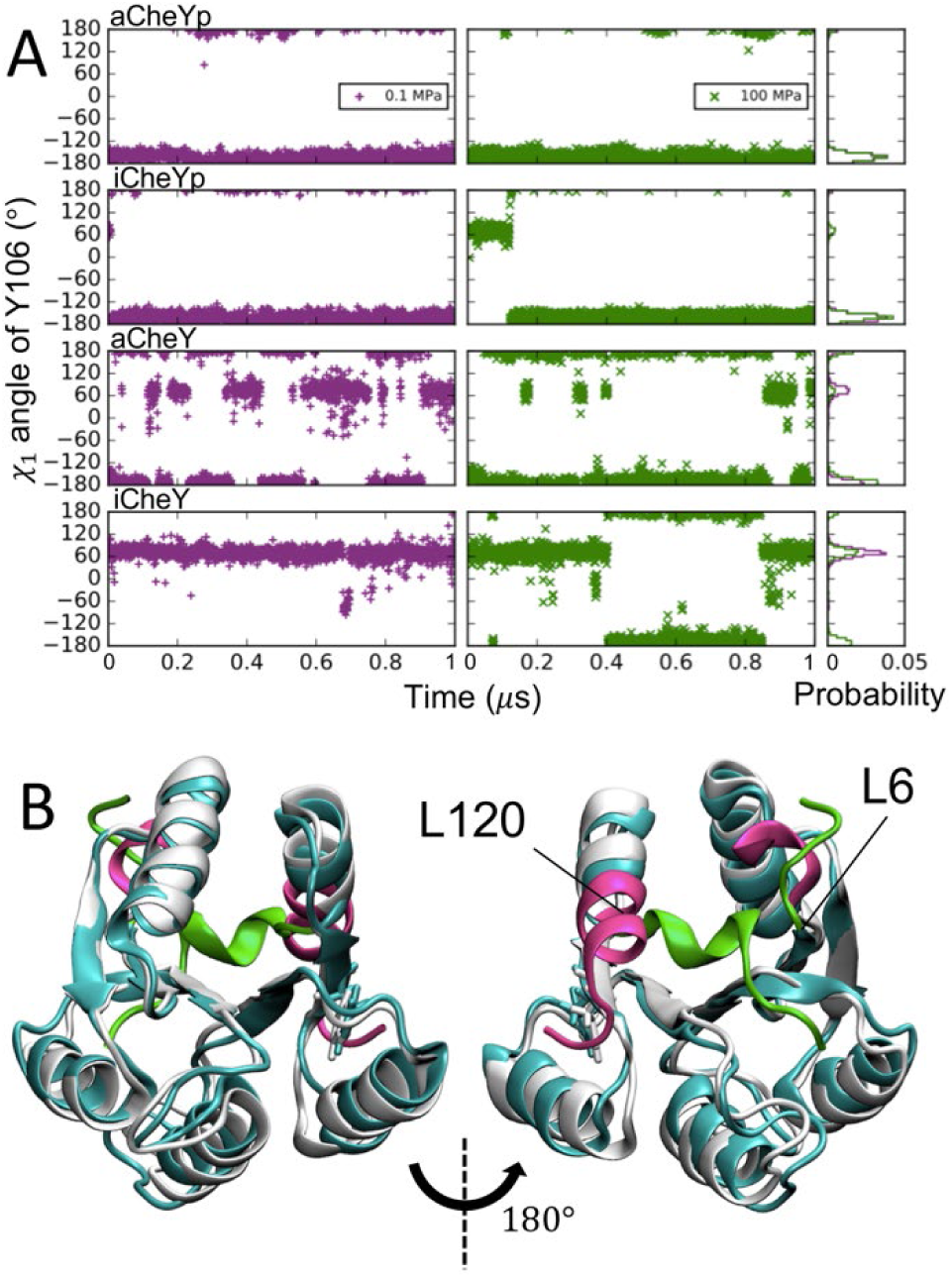
Pressure effects on the CheY monomer. **A.** Time evolution of the *χ*_1_ angle of Y106 during the 1-µs MD simulations for four different models of CheY at 0.1 (purple) and 100 MPa (green). Probability densities are shown on the right. **B.** Representative structures of aCheYp at 0.1 (white) and 100 MPa (cyan). A view from the opposite side is shown on the right. Segments significantly affected by pressure are shown in a different color, with the structures at 0.1 and 100 MPa indicated in magenta and green, respectively. Residues on the edges of the segments that underwent conformational transitions are labeled. The representative structures were determined as cluster centers by the k-means clustering of the last half of the MD trajectories.

### Pressure causes significant conformational change of CheY at 100 MPa

In the case of aCheYp at 100 MPa, we observed an interesting conformational transition of the N- and C-terminal segments (Fig. 2B); this transition began at approximately 0.5 μs and was completed at approximately 0.74 μs. The N-terminal conformational change involved the first five residues of CheY, a region that is far from the FliMN-binding site; the C-terminal conformational change occurred as a significant bending of the helix caused by large mainchain dihedral changes in Lys119 and Leu120, as revealed by Dihedral Transition Analysis (DTA)^66^. Since CheY residues 122, 123, and 126 are part of the interface residues of the complex, the binding affinity may be weakened by loss of some interactions if this conformational transition occurs upon binding; however, the bending occurred in the direction opposite to the FliM_N_-binding site (compare Fig. 1A and 2B), and this conformation is not expected to sterically interfere with FliM_N_ binding. In iCheY, a similar N-terminal change in the first five residues was observed at 100 MPa. In addition, we also identified a “sidechain flip”^66^ of Asp74, situated in at the end of the α-helix and far from the FliM_N_-binding interface.

### Pressure induces earlier detachment of FliM_N_

Next, we investigated pressure effects on the aCheYp–FliM_N_ complex by 1-μs MD at 0.1, 50, and 100 MPa. During the 1-μs MD, the complex structure was stable and its dissociation was not observed. Experimentally, dissociation of some protein complexes has been shown to occur at ∼ 60 MPa^14,67^, but simulating the dissociation process using standard MD is very difficult because the time scale of the complex dissociation can be much longer than the MD time scale^26^. Some of the residues between residues 83–125 of CheY make contacts with FliM_N_. The complex structure is stabilized by a salt bridge between Lys119 of CheY and Asp12 of FliM_N_ (Fig. S1) that was maintained during 90% of the simulation time, and also by a few mainchain–sidechain hydrogen bonds. Since the aforementioned conformational transition at 100 MPa starts from Lys119, that change may weaken the binding if the transition occurs. However, such a conformational change was not observed in the simulation of the complex at high pressure nor in the following dissociation simulations.

The dissociation process of the aCheYp and FliM_N_ complex was observed by PaCS-MD at 0.1, 50, and 100 MPa. In the present work, we focused on examining the pressure dependence of the binding affinity; five PaCS-MD trials were sufficient for this purpose, as will be shown below. In all the trials, dissociation up to 50 Å in the inter-center of mass distance, *d*, was observed within 400 cycles (Fig. 3A). This computational cost corresponds to a total simulation time of 40 ns (0.1 ns MD per replica × 400 cycles) and total computational cost of 400 ns (40 ns × 10 replicas). Relatively large variations in the number of PaCS-MD cycles required to cause the dissociation were consistent with the results of the earlier paper^31^, which showed that this number varied in different trials when 10 replicas were used. The generated dissociation pathways are shown in Figs. 3B and 3C. Although the direction of dissociation varied, the helical segment of the C-terminal segment of FliM_N_ always detached from aCheYp first, with subsequent unbinding of the N-terminal region (Fig. 3C-E). The former step, consisting of detachment of the FliM_N_ C-terminal helix, occurred at *d* = 25 ± 2, 25 ± 2, and 27 ± 2 Å at 0.1, 50, and 100 MPa (mean ± standard deviation (SD)), respectively, indicating a lack of clear pressure dependence. This process correlated with breaking of the aforementioned salt bridge (Lys119: CheY–Asp12: FliM_N_), which occurred slightly earlier at *d* = 20 – 21 Å. The subsequent unbinding of the N-terminal region occurred at *d* = 45 ± 3, 41 ± 32, and 35 ± 4 Å at 0.1, 50, and 100 MPa, respectively, suggesting that higher pressure induces earlier complete detachment of FliM_N_. We regard the phase before the helix detachment as the bound state (Phase 1), the phase before the complete dissociation as the partially bound state (Phase 2), and the last phase as the unbound state (Phase 3). This pressure-dependent dissociation is investigated further below.

**Figure 3.**
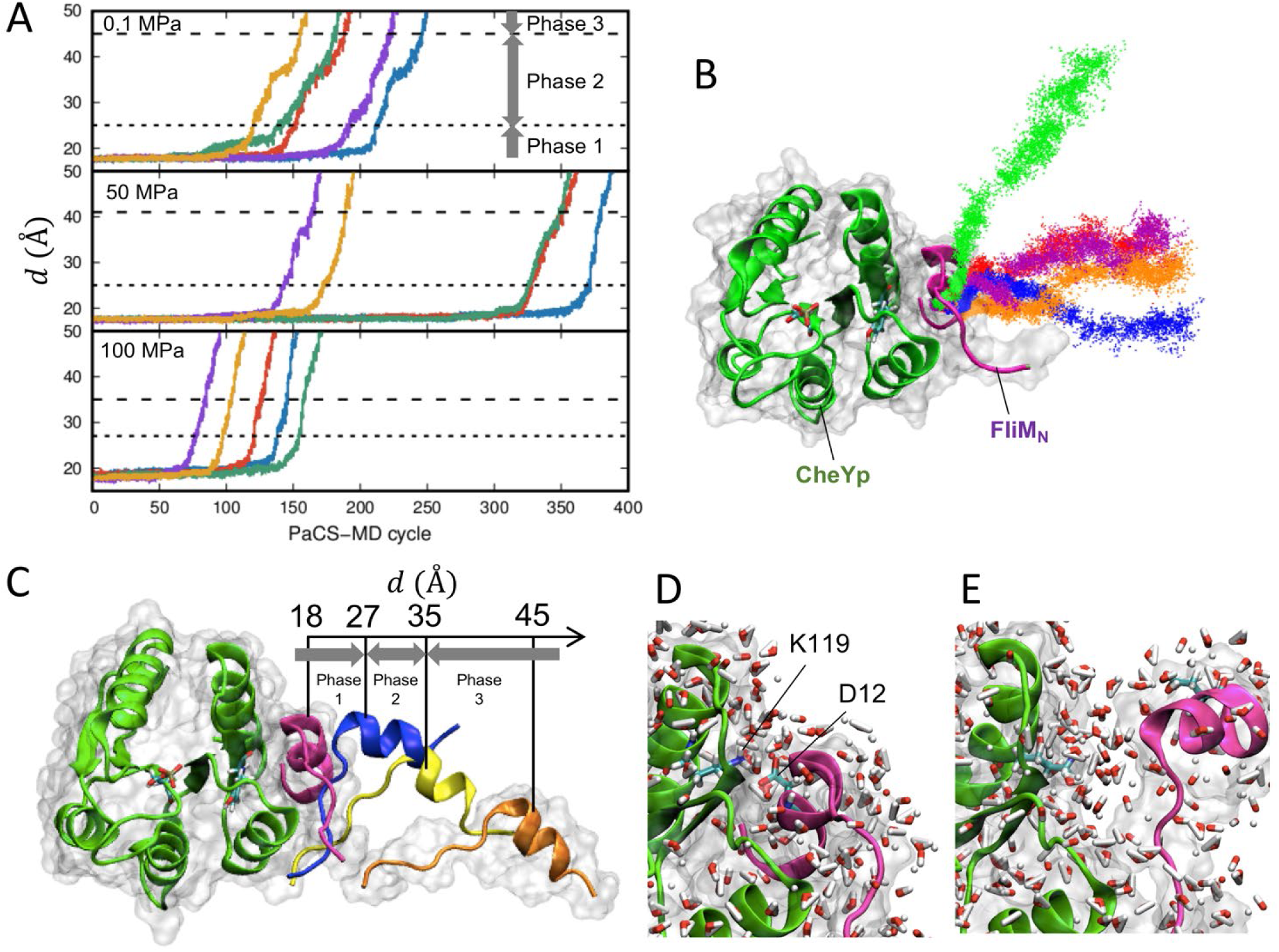
Dissociation of the CheYp–FliM_N_ complex at 0.1, 50, and 100 MPa, as simulated by PaCS-MD. **A.** The inter-COM distance between CheYp and FliM_N_, with *d* indicated as a function of the PaCS-MD cycle at 0.1 (top), 50 (center), and 100 (bottom) MPa. Five independent PaCS-MD simulations are shown in different colors. The broken lines indicate the borders between Phases 1-2 and 2-3. **B.** COM positions of FliM_N_ obtained by PaCS-MD and all additional MD simulations at 100 MPa. Color differences denote different trials. **C.** Examples of snapshots of FliM_N_ during a dissociation at 100 MPa. MD snapshots with four different *d* values are shown in different colors after superimposing CheYp. Molecular surfaces of the initial and last snapshots are shown in transparent colors. **D, E.** Closeup view of the interface, **D** before the breakage of the key salt bridge (Lys119: CheY–Asp12: FliM_N_) at *d* = 18 Å, and **E** just after the detachment of the FliM_N_ helix at *d* = 27 Å at 100 MPa. Water molecules in the first hydration shell of each protein are shown with stick models and atomic-color basis.

In addition, we examined pressure effects on monomeric FliM_N_ by analyzing trajectories generated by two different procedures. In the first procedure, MD simulations were started from the FliM_N_ structure taken from the crystal structure of the complex (PDB ID: 1F4V^36^) and solvated as a monomer. MDs lasting 0.5 μs were conducted at 0.1, 50, and 100 MPa and analyzed. In the second procedure, the FliM_N_ structures completely dissociated from aCheYp (*d* > 40 Å) by PaCS-MD were analyzed. FliM_N_ in the former tended to maintain the helical region formed in the CheY-bound state, whereas in the latter, FliM_N_ was slightly more unstructured in the helical region, a change that was induced during the detachment process.

### Thermodynamic properties of the proteins are conserved up to 100 MPa

Various properties of FliM_N_ and aCheYp in monomeric and complex states were calculated at different pressures (Table S1). The C_α_ root-mean-square deviation (RMSD) of the MD-representative structure at high pressures compared to that at 0.1 MPa was small (∼ 1 Å) for monomeric CheY and the aCheYp–FliM_N_ complex, except for monomeric aCheYp and iCheY at 100 MPa, in which the aforementioned conformational changes occurred. The RMSD of monomeric FliM_N_ is larger, mainly due to the structural fluctuation of the unstructured segment at the N-terminus, which is also very flexible in the complex form. In the helical region of FliM_N_ (residues 8–15), the RMSD was much smaller. These results indicated that hydrostatic pressures up to ∼ 50 MPa do not significantly change the structures of the monomers and complex within the range of the simulations, but suggested that 100-MPa pressure can change the structure of monomeric CheY. SASA, *V*_ex_, cavity volume (*V*_cav_), and *κ*_*T*_ also showed no significant change between the two pressure conditions (Table S1). It is worth mentioning that the TIP3P model^41^ used in the current simulation has been reported to reproduce experimental compressibility up to 200 MPa^68^. Since CheY is a small protein, the *V*_cav_ of CheY was very small even at 0.1 MPa and shrank slightly at 100 MPa; however, the amount of shrinkage was equivalent only to the size of one water molecule, which is smaller than the observed cavity and volume fluctuations. The pressure-independent behavior of *k*_*T*_ is consistent with the fact that protein structure does not significantly change in this pressure range, indicating that high pressure mainly compresses water. This result is considered to be reasonable, given that the *κ*_*T*_ of bulk water is ∼ 2-fold larger than that of typical proteins^10,69^. This observation also is consistent with the Young’s moduli of hydrated proteins, which are in the range of 1 ∼ 3 GPa for actin, tubulin, and flagellin^70^. The Young’s modulus of CheY should be equal to or greater than those of these structural protein, given that CheY is a smaller and more-compact globular protein. Expected elastic deformation at 100 MPa should be 3.3% if Young’s modulus is 3 GPa, which suggests very small deformation comparable to thermal fluctuations (∼ 1 Å).

### Pressure increases protein hydration and induces water penetration into the interface

Next, we analyzed the hydrogen bond network, including water molecules. The applied pressure did not alter the number of hydrogen bonds connecting protein-protein (*H*_PP_) that sums up both intra- and inter-protein hydrogen bonds (Table S1). The result is consistent with the aforementioned result that high pressure ≤ 100 MPa does not change protein structure in most cases. Similarly, the number of protein-water-protein hydrogen bond network (*H*_PWP_) was almost pressure-invariant. On the other hand, the number of protein-water hydrogen bonds (*H*_PW_) in monomeric CheY and the complex increased as the pressure increased, which agrees with the expectation that high pressure results primarily in the compression of water. Similar changes in the hydrogen bonding network at the protein-water interface previously have been reported to be induced by pressure^23^.

Next, we investigated water molecules in the first and second hydration shells around the proteins, namely the first and second shell surface waters, which are defined as the water molecules within 3 and 3–6 Å (respectively) from any protein atoms. We counted the number of surface waters in the first and second shells, *N*_1SW_ and *N*_2SW_, respectively. Interestingly, significant increases in hydrated water molecules at high pressure were observed (Fig. 4A), consistent with the aforementioned increase in *H*_PW_ (Table S1). Compared to 0.1 MPa, ∼ 30 more water molecules were found in the first hydration shell of monomeric CheY and the aCheYp–FliM_N_ complex at 100 MPa, which is equivalent to ∼ 8% increase in the ratio (*r*_1SW_). This increase was slightly higher than the average increase in the density of bulk water at the increased pressure (∼ 5%). Bulk water densities at 0.1, 50, and 100 MPa were calculated to be 3.29 ± 0.70, 3.37 ± 0.69, and 3.45 ± 0.67 × 10^−2^ Å^-3^, respectively (see Methods for detail). These results indicated that the pressure increase induces an increase of water density around the protein surfaces compared to that of bulk water. This observation is consistent with previous theoretical and experimental studies of proteins in solution at similar pressures^1,3,11,21^. This tendency of hydration increase at higher pressure also was seen in the second hydration shell (∼ 6% increase in *r*_2SW_) but the increase was nominally less than that seen in *r*_1SW_. The increase of hydration water was exceptionally higher for aCheYp in the monomernic state at 100 MPa, in which (as mentioned earlier) a conformational transition of the C-terminal helix was seen. The same tendency also was observed in the number of interface waters, which was analyzed as described below. This observation implied that the conformational transition promotes greater hydration of aCheYp at higher pressure.

**Figure 4.**
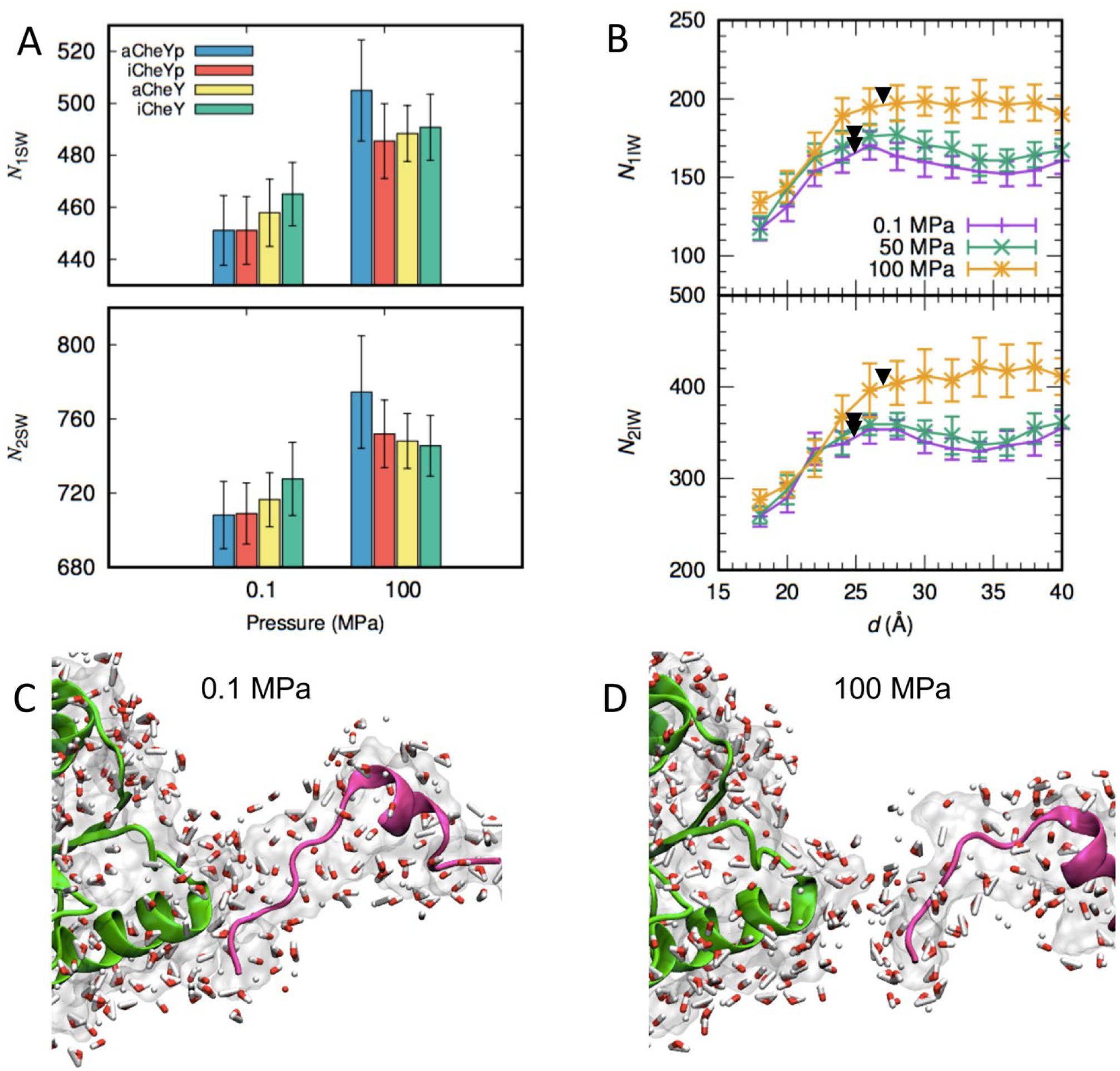
Pressure effects on protein hydration. **A**. Pressure dependence of the number of surface waters in the first and second shells, *N*_1SW_ (upper panel) and *N*_2SW_ (lower panel), respectively. **B.** Change in the numbers of interface waters, *N*_1IW_ (upper panel) and *N*_2IW_ (lower panel), during the dissociation process along *d* at 0.1 (purple), 50 (green), and 100 MPa (orange). The error bars show standard deviations among the five dissociation simulations. Black triangles indicate the borders between Phases 1 and 2. **C, D.** MD snapshots at *d* = 40 Å at **C** 0.1 or **D** 100 MPa. Molecular representation is the same as in Fig. 3C and D.

We further focused on hydration around the complex interface and conducted similar analysis of the number of interface water in the first (*N*_1IW_) and second (*N*_2IW_) hydration shells around the interface residues (Table S1). These residues are defined as the residues involved in CheYp-FliM_N_ contacts during more than 80% of the time of the second half of the 1-*μ*s MD trajectory at 0.1 MPa. As expected, the increase in *N*_1IW_ was significant in all cases. For example, the water density of the interface in the first shell in the complex increased 6% when comparing between 0.1 MPa and 50 MPa, and 8% when comparing between 0.1 MPa and 100 MPa (see *r*_1IW_ in Table S1), while those of the second shell increased 5% and 4%, respectively (see *r*_2IW_ in Table S1). This result also showed that, in both the protein surface and protein-protein interface, the first shell water density tends to increase more than that of the second shell. High pressure increases the water density of the first hydration shell around the proteins and provokes water penetration into the interface. As shown above, high pressure did not change the timing of the salt bridge breakage, which implied that high pressure has a relatively weak effect on the salt bridge stability. Once salt bridge breakage occurred in the initial stage of dissociation, enhanced water penetration at high pressure accelerated the complete dissociation.

During the dissociation process of the complex, *N*_1IW_ and *N*_2IW_ gradually increased until the detachment of the C-terminal helix of aCheYp at the end of Phase 1, which is indicated by triangles in Fig. 4B. At this moment, the number of interface waters at 100 MPa was significantly greater than those at 0.1 and 50 MPa, which indicates the nonlinearity of the high-pressure effect on hydration at the protein-protein interface. As noted above, complete dissociation occurred at shorter *d* as pressure increased. This acceleration was caused by the increased hydration at higher pressure (Fig. 4C and D). At 0.1 MPa and *d* = 40 Å, the complex was still in Phase 2 (Fig. 4C) but completely dissociated at the same distance at 100 MPa because more water molecules penetrated into the complex interface (Fig. 4D).

### Pressure impairs CheYp–FliMN binding

The free energy profile (potential of mean force as a function of *d*) from the bound to unbound states of CheYp and FliM_N_ at 0.1, 50, and 100 MPa is shown in Fig. 5A. This showed that the free energy difference between the bound and unbound states significantly decreases as the pressure increases, which clearly indicates weaker binding affinity at higher pressure. Although only five PaCS-MD trials were conducted, the free energy profiles showed small standard deviations, indicating that free energy converges to a certain value independent of dissociation direction if two molecules are sufficiently separated and interactions between the molecules become negligibly small, as shown in earlier work^29,31^. It should be noted that the free energy converged upon complete detachment of the complex, shown by red triangles in Fig. 5A. Since free energy differences among different pressures in Phase 1 (up to red open triangles) also were relatively small, the overall free energy differences originated mainly from the changes that occurred in Phase 2. From the point of view of the free energy profile, this difference may be interpreted as the result of weakened long-range interactions between the two proteins by more hydrated waters infiltrating between the proteins at higher pressure.

**Figure 5.**
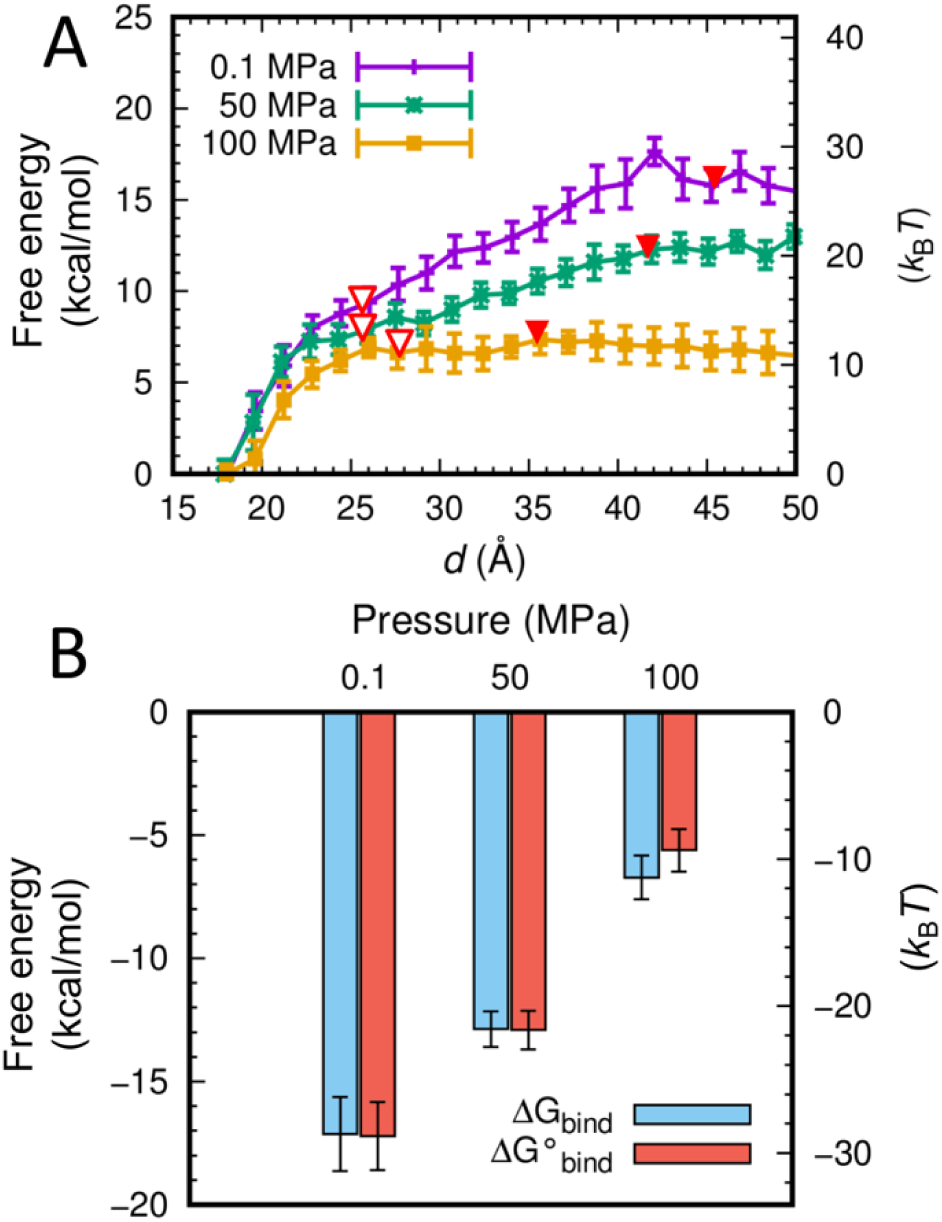
Pressure effects on binding free energy. **A.** Dissociation free energy profiles as a function of *d* calculated by PaCS-MD/MSM. Red open and filled triangles indicate the borders between Phases 1-2 and 2-3, respectively. **B.** Standard binding free energies calculated from the free energy profiles and volume correction 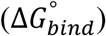 and binding free energies before volume correction (*ΔG*_*bind*_).

The standard binding free energy (*ΔG°*_bind_) of aCheYp and FliM_N_ was calculated from the free energy profile (see Methods) and is shown in Fig. 5B. The *ΔG°*_bind_ value significantly decreased with an increase of hydrostatic pressure. This observation is qualitatively consistent with the model that protein-protein binding is suppressed by hydration of protein surfaces induced by high hydrostatic pressure^1^. In the case of the CheYp–FliM_N_ complex, *ΔG°*_bind_ values were −17.2 ± 1.4, - 12.9 ± 0.8, and −5.6 ± 0.9 kcal/mol at 0.1, 50, and 100 MPa, respectively, indicating that high pressures up to 100 MPa weakened binding between the two proteins. The change in the *ΔG°*_bind_ value from 50 to 100 MPa (−7.3 kcal/mol) was greater than that from 0.1 to 50 MPa (−4.3 kcal/mol), which is consistent with the non-linear pressure response shown above. Overall, the simulation results suggested that the binding affinity of the CheYp protein to the flagellar motor protein FliM is significantly lowered by high pressure.

### Pressure inhibits CW rotation of *E. coli* flagellar motors

Finally, we experimentally examined the effect of hydrostatic pressure on the rotation of the flagellar motor in *E. coli* strain SYC12 (wild type for chemotaxis). Rotation of a single motor was monitored as rotation of a cell body by attaching a single flagellar filament extended from the cell body directly to an inserted coverslip in the high-pressure chamber (Fig. 6A). Figure 6B shows typical time courses of the rotational speed of a single motor at 0.1 and 40 MPa, respectively. At 0.1 MPa, the flagellar motor rotated smoothly, and the rotational direction frequently switched from CCW to CW, and vice versa. In contrast, the motor rotated exclusively in a CCW direction at 40 MPa. The CW bias values, the time occupancy of CW rotation, at 0.1 and 40 MPa were 0.19 and zero, respectively (*n*=25 cells) (Fig. 6C). On the other hand, application of 40 MPa showed no significant effect on the rotational speed (Fig. 6D). After the release of pressure, the frequent switching of the motor was observed to resume. These results indicated that application of 40 MPa of pressure suppresses motor switching, without affecting the overall structure and function of the motor. Therefore, the application of pressure is suggested to inhibit the binding of CheYp to FliM in *E. coli* cells, which is consistent with our observations using the simulations.

**Figure 6.**
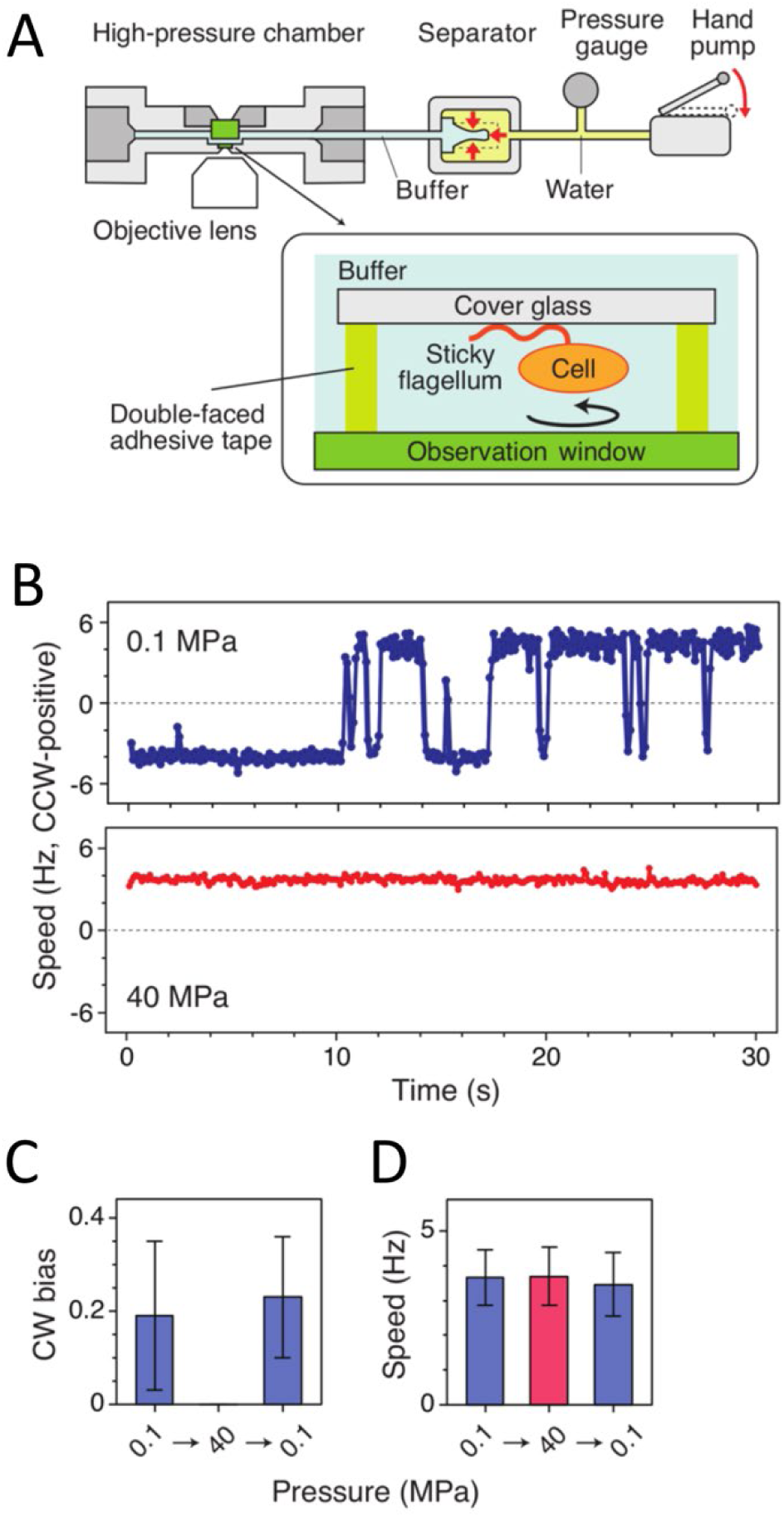
Rotation of flagellar motors at high pressure. **A.** Schematic illustration of the experimental setup. **B.** Time courses of the rotational speed of the same motor of the *E. coli* strain (wild type for chemotaxis) at 0.1 (top) and 40 MPa (bottom). **C** and **D.** Pressure response of the rotational directionality (C) and speed (D). The rotation of the same motors was tracked when the pressure was changed from 0.1 to 40 MPa, and then returned to 0.1 MPa (mean ± SD, *n*=25).

## Conclusion

In this work, we first showed, by MD simulation, that the binding of chemotaxis signaling protein CheYp to motor rotor protein FliM_N_ is inhibited at high pressure. Even at 100 MPa, the active form of CheYp was shown to retain its overall structure at ambient pressure, but high pressure increased the water density in the first hydration shell and caused conformational change of the C-terminal helix in the monomeric aCheYp case. Using dissociation simulation of the CheYp–FliM_N_ complex by PaCS-MD and subsequent MSM analysis, we demonstrated that the binding affinity of the two proteins weakens as pressure rises, as shown by the calculated standard binding free energy. Consistent with these results, we showed, by high-pressure microscopy, that high pressure reversibly suppresses CW rotation of the bacterial flagellar motor. Altogether, our results provide a clear picture of high-pressure inhibition of CheYp–FliM_N_ binding.

To the best of our knowledge, this is the first molecular simulation that observed the pressure-induced dissociation process of protein complexes in solution without applying bias. Since the calculation of binding free energy between proteins requires very long computational time, the pressure dependence of the binding free energy between proteins previously has been argued based on calculations for smaller methane-like molecules^71,72^. According to the heteropolymer collapse theory, hydrophobic interactions are destabilized by pressure ≫ 100 MPa, where the pressure-induced change in the interfacial free energy between protein and water lead to the penetration of water into a protein’s hydrophobic core and result in protein denaturation^73^. In the present work, we demonstrated that the binding free energy between protein molecules decreases as pressure increases in the range ≤ 100 MPa. High pressure in this range induces a significant increase of water density around proteins; this increase is higher than that of bulk water density, especially in the first hydration shell, which apparently facilitates water penetration into the protein complex interface. We also observed a breakage of a key salt bridge accompanied by dissociation of the protein complex. Water molecules around such broken ionic bonds have been known to form a hydration shell that has a higher packing density than that of bulk water (i.e., electrostriction), which decreases the volume of a system and favors rupture of ionic bonds at high pressure^74^.

In conclusion, high pressure modulates a protein-protein interaction by changing its hydration state even at 100 MPa or below. This pressure range is equivalent to the pressure in the deep sea, although studies for pressure effects on proteins have focused primarily on the denaturation (or unfolding) of proteins caused by higher pressures^4,9,74^. Studies on the effects of such relatively high pressure on proteins in the biosphere are essential for understanding proteins’ bioresponsive functions at the molecular level and also for designing functions governed by protein-protein interactions.

## Acknowledgments

We thank Dr. Yong-Suk Che for providing *E. coli* strain SYC12. This research was supported in part by MEXT/JSPS KAKENHI grants to I.K., M.N., and A.K. (No. JP17KT0026), to A.K. (No. JP19H03191), to M.N. (Nos. JP16K04908 and JP17H05880), and to Y.S. (No. JP19H05404), and by a MEXT grant (“Priority Issue on Post-K Computer” (Building Innovative Drug Discovery Infrastructure Through Functional Control of Biomolecular Systems)) to A.K. The computations were performed in part using the supercomputers at The Research Center for Computational Science (RCCS), The National Institute of Natural Science, and at The Institute of Solid State Physics (ISSP), The University of Tokyo. This research also used computational resources of the K computer provided by the RIKEN Advanced Institute for Computational Science through the HPCI System Research project (Project IDs: hp150270, hp160207, hp170254, hp180201, and hp190181).

## Supplementary Material

A table contains various properties of protein structures, a visualization of the residues mentioned in the text, and a photograph of the experimental setup.

