## Supplemental Materials for "High pressure inhibits signaling protein binding to the flagellar motor and bacterial chemotaxis through enhanced hydration"

*[1]School of Life Science and Technology, Tokyo Institute of Technology, Ookayama, 2-12-1 Meguro-ku, Tokyo, 152-8550, Japan, [2]Institute of Molecular and Cellular Biosciences, The University of Tokyo, 1-1-1 Yayoi, Bunkyo-ku, Tokyo 113-0032, Japan,[3]Department of Physics, Kindai University, 3-4-1 Kowakae, Higashiosaka, Osaka 577-8502, Japan, [4]Department of Frontier Biosciences, Hosei Univesity, Koganei, Tokyo 184-0002, Japan*

*

**Table S1.** Effect of high pressure on the CheY and FliM$_N$ structures and various properties.

| Model | State[a] | Pressure [MPa] | RMSD[b] [Å] | SASA[c] [$10^3$ Å$^2$] | $V_{ex}$ [d] [$10^4$ Å$^3$] | $V_{cav}$ [e] [Å$^3$] | $\kappa$[ef] [GPa$^{-1}$] | $H_{PP}$[g] | $H_{PW}$[h] | $H_{PWP}$[i] | $r_{1SW}$[j] | $r_{1IW}$[k] | $r_{2SW}$[k] | $r_{2IW}$[m] |
|---|---|---|---|---|---|---|---|---|---|---|---|---|---|---|
| aCheY | M | 0.1 | - | 7.1 ± 0.1 | 2.41 ± 0.02 | 6 ± 5 | 0.2 ± 0.3 | 57 ± 6 | 208 ± 10 | 18 ± 3 | - | - | - | - |
| | | 100 | 0.9 | 7.1 ± 0.1 | 2.40 ± 0.01 | 3 ± 3 | 0.2 ± 0.2 | 57 ± 6 | 215 ± 10 | 19 ± 4 | 1.07±0.04 (31 ± 17) | 1.03 ± 0.08 (3 ± 8) | 1.04 ± 0.03 (32 ± 21) | 1.03 ± 0.07 (6 ± 17) |
| iCheY | M | 0.1 | - | 7.2 ± 0.2 | 2.43 ± 0.02 | 6 ± 5 | 0.3 ± 0.4 | 57 ± 6 | 210 ± 11 | 18 ± 4 | - | - | - | - |
| | | 100 | 2.0 | 7.1 ± 0.1 | 2.41 ± 0.01 | 5 ± 5 | 0.2 ± 0.3 | 54 ± 6 | 221 ± 9 | 21 ± 4 | 1.06 ± 0.04 (26 ± 18) | 1.07 ± 0.11 (7 ± 11) | 1.02 ± 0.04 (18 ± 26) | 1.00 ± 0.07 (1 ± 17) |
| aCheYp | M | 0.1 | - | 7.0 ± 0.1 | 2.41 ± 0.01 | 8 ± 5 | 0.2 ± 0.3 | 58 ± 5 | 208 ± 9 | 20 ± 4 | - | - | - | - |
| | | 100 | 2.9 | 7.2 ± 0.3 | 2.41 ± 0.03 | 6 ± 4 | 0.7 ± 0.7 | 60 ± 6 | 211 ± 11 | 19 ± 4 | 1.12 ± 0.05 (54 ± 24) | 1.22 ± 0.12 (21 ± 10) | 1.09 ± 0.05 (66 ± 35) | 1.17 ± 0.11 (39 ± 24) |
| | D | 0.1 | - | 7.0 ± 0.1 | 2.41 ± 0.0 | 5 ± 2 | 0.2 ± 0.3 | 58 ± 5 | 211 ± 1[illegible] | 21 ± 4 | - | - | - | - |
| | | 50 | 1.2 | 7.0 ± 0.1 | 2.40 ± 0.0 | 4 ± 2 | 0.1 ± 0.2 | 58 ± 5 | 216 ± 8 | 19 ± 3 | 1.05 ± 0.04 (23 ± 17) | 1.10 ± 0.09 (9 ± 8) | 1.04 ± 003 (28 ± 22) | 1.05 ± 0.06 (10 ± 13) |
| | | 100 | 1.2 | 7.1 ± 0.1 | 2.39 ± 0.0 | 3 ± 4 | 0.1 ± 0.2 | 58 ± 5 | 214 ± 8 | 18 ± 3 | 1.08 ± 0.04 (34 ± 17) | 1.16 ± 0.09 (15 ± 8) | 1.06 ± 0.03 (42 ± 22) | 1.10 ± 0.06 (22 ± 14) |
| iCheYp | M | 0.1 | - | 7.0 ± 0.1 | 2.40 ± 0.01 | 9 ± 6 | 0.2 ± 0.3 | 59 ± 5 | 207 ± 8 | 19 ± 4 | - | - | - | - |
| | | 100 | 0.9 | 7.1 ± 0.2 | 2.40 ± 0.02 | 7 ± 6 | 0.3 ± 0.4 | 59 ± 5 | 215 ± 10 | 20 ± 4 | 1.08 ± 0.04 (34 ± 19) | 1.09 ± 0.10 (9 ± 9) | 1.06 ± 0.04 (43 ± 25) | 1.04 ± 0.08 (8 ± 18) |
| FliM$_N$ | M | 0.1 | - | 1.7 ± 0.1 | 0.35 ± 0.01 | 0 | 0.5 ± 0.6 | 3 ± 2 | 39 ± 4 | 2 ± 1 | - | - | - | - |
| | | 50 | 0.5 | 1.7 ± 0.1 | 0.35 ± 0.01 | 0 | 0.6 ± 0.8 | 4 ± 2 | 39 ± 4 | 3 ± 1 | 1.07 ± 0.13 (7 ± 14) | 1.06 ± 0.15 (5 ± 12) | 1.05 ± 0.11 (14 ± 28) | 1.06 ± 0.12 (12 ± 25) |
| | | 100 | 0.3 | 1.7 ± 0.1 | 0.35 ± 0.01 | 0 | 0.3 ± 0.4 | 4 ± 2 | 39 ± 4 | 2 ± 1 | 1.09 ± 0.12 (11 ± 13) | 1.08 ± 0.14 (7 ± 12) | 1.07 ± 0.11 (17 ± 26) | 1.04 ± 0.11 (8 ± 23) |
| | D | 0.1 | - | 2.0 ± 0.1 | 0.37 ± 0.0 | 0 | 0.2 ± 0.3 | 2 ± 1 | 31 ± 4 | 1 ± 1 | - | - | - | - |
| | | 50 | 1.1 | 1.9 ± 0.2 | 0.36 ± 0.0 | 0 | 0.9 ± 1.2 | 4 ± 2 | 28 ± 4 | 1 ± 1 | 1.11 ± 0.07 (12 ± 7) | 1.11 ± 0.09 (9 ± 7) | 1.21 ± 0.09 (89 ± 33) | 1.20 ± 0.10 (65 ± 28) |
| | | 100 | 0.6 | 1.9 ± 0.1 | 0.36 ± 0.0 | 0 | 0.2 ± 0.2 | 3 ± 1 | 29 ± 5 | 1 ± 1 | 1.00 ± 0.06 (0 ± 7) | 1.01 ± 0.08 (0 ± 6) | 1.01 ± 0.07 (6 ± 28) | 1.03 ± 0.08 (9 ± 25) |
| aCheYp–FliM$_N$ | C | 0.1 | - | 7.3 ± 0.1 | 2.64 ± 0.01 | 6 ± 6 | 0.2 ± 0.3 | 73 ± 6 (6 ± 2) | 229 ± 10 | 22 ± 5 (2 ± 1) | - | - | - | - |
| | | 50 | 1.1 | 7.5 ± 0.2 | 2.66 ± 0.01 | 8 ± 4 | 0.2 ± 0.3 | 72 ± 7 (5 ± 2) | 234 ± 11 | 23 ± 4 (2 ± 1) | 1.06±0.04 (27 ± 20) | 1.06±0.09 (7 ± 9) | 1.04±0.04 (30 ± 29) | 1.05±0.06 (12 ± 16) |
| | | 100 | 1.1 | 7.5 ± 0.1 | 2.64 ± 0.02 | 4 ± 2 | 0.2 ± 0.3 | 73 ± 6 (6 ± 2) | 239 ± 11 | 22 ± 4 (2 ± 1) | 1.08±0.04 (36 ± 18) | 1.08±0.09 (8 ± 10) | 1.06±0.04 (46 ± 27) | 1.04±0.06 (11 ± 14) |

[a]M: Monomer state started from monomeric structure, D: Dissociated state generated by PaCS-MD, C: Complex state. The average values and the standard deviation (values after '±') for the second half of the MD trajectories (M and C), and those for the PaCS-MD trajectories in Phase 3 (D) are shown. [b]RMSD of the representative structure at 100 MPa from that at 0.1 MPa calculated for all $C_\alpha$ atoms except for two terminal residues. [c]Solvent accessible surface area. [d]Excluded volume defined as the space inside of SASA. [e]Cavity volume inside the proteins. [f]Isothermal compressibility. [g-i]$H_{PP}$, $H_{PW}$, and $H_{PWP}$ are the number of hydrogen bonds connecting protein–protein, protein–water, and protein–water–protein. For $H_{PP}$, the values in the parentheses indicate the number between aCheYp and FliM$_N$. For $H_{PWP}$, the values in the parentheses show the number for aCheYp–water–FliM$_N$. [j]Ratio of the number of water molecules in the first hydration sphere relative to 0.1 MPa. [k]Ratio of the number of water molecules around the complex interface in the first hydration sphere relative to 0.1 MPa. [l]Ratio of the number of water molecules in the second hydration sphere relative to 0.1 MPa. [m]Ratio of the number of water molecules around the complex interface in the second hydration sphere relative to 0.1 MPa. [j-m]The values in the parentheses indicates actual increase in the number of waters.

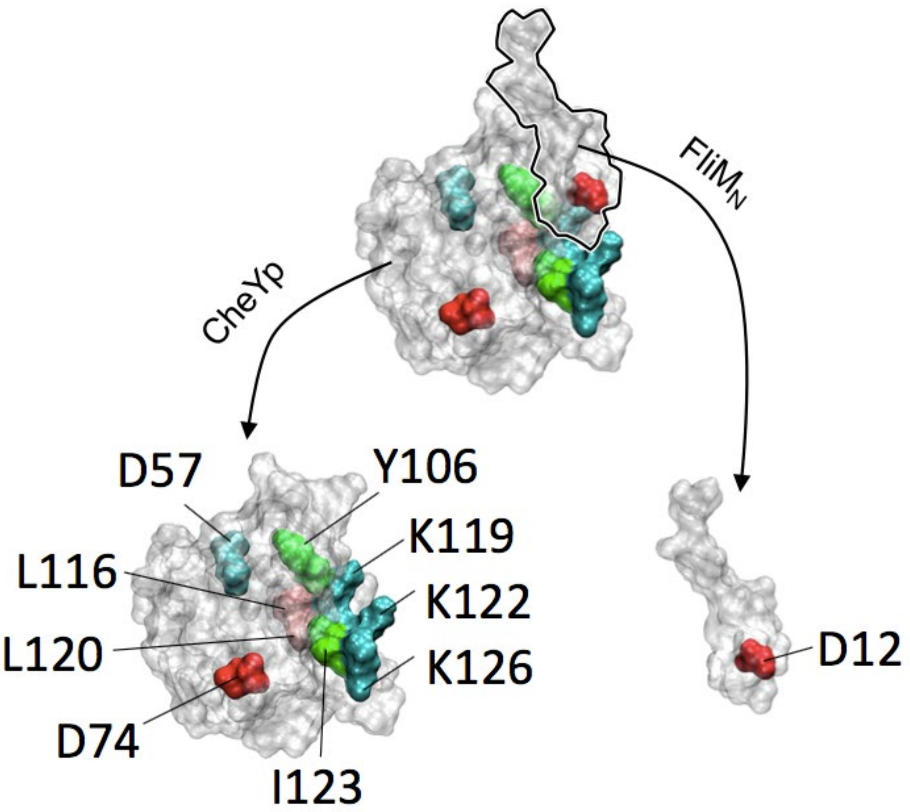


**Figure S1.** The residues mentioned in the main text. Phosphorylated D57, interface residues (Y106 –K126 of CheY and D12 of FliM$_N$), and D74 that underwent side-chain flip at 100 MPa in iCheY are shown. These residues are colored by residue types and the other residues are shown in transparent white so that residues in the back side are visible, e.g., D12.

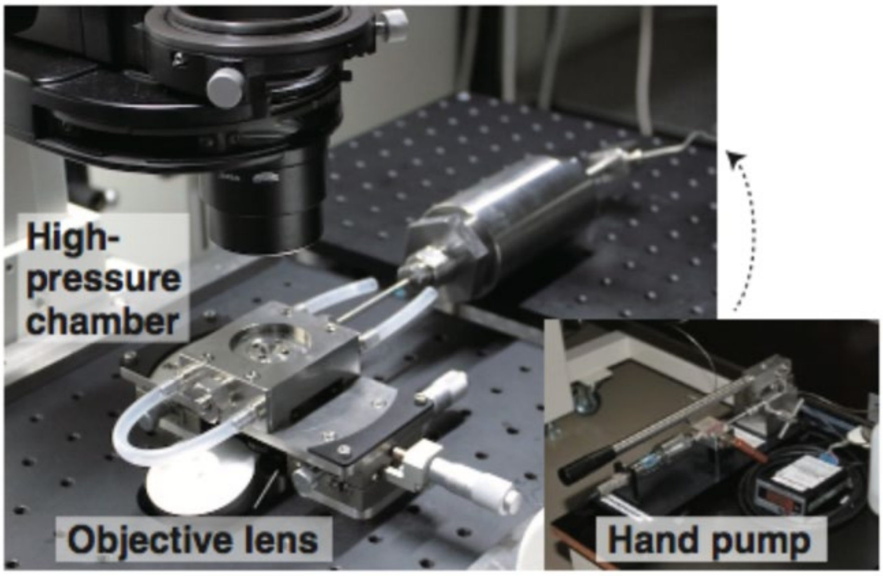


**Figure S2.** Photograph of the experimental setup.
